# Generating closed bacterial genomes from long-read nanopore sequencing of microbiomes

**DOI:** 10.1101/489641

**Authors:** Eli L. Moss, Ami S. Bhatt

**Affiliations:** Department of Medicine (Hematology, Blood and Marrow Transplantation) and Department of Genetics, Stanford University, Stanford, California, USA

## Abstract

We present the first method for efficient recovery of complete, closed genomes directly from microbiomes using nanopore long-read sequencing and assembly. We apply our approach to three healthy human gut communities and compare results to short read and read cloud approaches. We obtain nine finished genomes including the first reported closed genome of *Prevotella copri*, an organism with highly repetitive genome structure prevalent in non-western human gut microbiomes.

## Main Text

*De novo* reconstruction of complete microbial genomes from metagenomes has been a longstanding goal of microbiome research. Although current reference-based methods are able to detect known organisms and genes in metagenomes, only *de novo* approaches are able to characterize novel genome sequences, or accurately place mobile or transferred elements in new genomic contexts. The tremendous diversity and plasticity of bacterial genomes, as well as the difficulty of bacterial isolation and culture, demand effective culture-free methods for producing genomes directly from metagenomes.

Current metagenomic sequencing and assembly methods do not typically yield finished bacterial genomes, although previous efforts have achieved single closed genomes in simple communities^1^, or multiple genomes with skilled manual assembly and scaffolding^2^. Consequently, genome drafts are formed by grouping (i.e. binning) similar contigs within fragmentary assemblies^3,4^. This is an imperfect process, often compromising the purity or the completeness of the genome reconstruction. As assembly contiguity increases, the sensitivity and specificity of genome binning are improved as fewer, larger contigs need to be grouped to form each genome. Indeed, at the point when genomes are assembled in single contigs, binning becomes unnecessary. Nanopore long read assembly has yielded complete genomes in cultured bacterial isolates^5–8^, suggesting potential for effective assembly in more complex microbial communities. However, the performance of nanopore and other long read approaches in metagenomic sequencing and assembly has been limited by the lack of effective and efficient methods to maximize molecular weight, mass yield and purity of DNA extracted from these samples.

We present a workflow consisting of stool DNA extraction, nanopore sequencing, assembly and post-processing steps capable of producing multiple complete, circular bacterial genomes directly from metagenomes. Our extraction approach produces microgram quantities of pure, high molecular weight (HMW) DNA suitable for long read sequencing from as little as 300 milligrams (mg) of stool. Our computational workflow, consisting of assembly and post-processing, does not involve manual intervention in assembly, scaffolding, bacterial isolation, or existing reference coverage of the target metagenome. Thus, this workflow is the first to provide a rapid, simple, cost-effective, automated approach to close high numbers of bacterial genomes directly from metagenomic samples.

Short read and read cloud data and assemblies for samples P1 and P2-A were used without modification as previously described^9^. The standard approach used to extract DNA for these libraries produced fragmented DNA which incurred a severe loss with size selection, necessitating approximately 300 mg input stool to assure the 1 nanogram (ng) final HMW DNA mass required for read cloud library preparation. Current long read library prep protocols require 1000 ng of HMW input DNA, well beyond the practical capability of existing stool DNA extraction techniques. In order to maximize the throughput and read length of nanopore sequencing, a new approach yielding DNA in dramatically higher quantity and molecular weight was needed.

We developed a method for HMW extraction capable of yielding 1000-fold more DNA over 5 kb than a conventional bead-beating approach (Supplementary Figure 1, see Methods). We applied this method to two samples (P1 and P2-A) as well as a third sample (P2-B), collected 15 months later from the second individual. HMW DNA extraction yielded at least 1 μg HMW DNA per 300 mg input stool mass for all samples (Supplementary Table 1). Nanodrop measurement produced A_260/280_ ratios over 1.86 and A_260/230_ ratios over 2.23 for all samples, indicating absence of contaminants such as proteins, solvents and salts.

We obtained a total of 12.7 giga-base pairs (Gbp), 6.1 Gbp, and 7.6 Gbp of long read data for samples P1, P2-A, and P2-B, respectively (Supplementary Figure 2, Supplementary Table 2) with N50 values of 4.7 kbp, 3 kbp and 3 kbp. The taxonomic composition of reads obtained through our approach was compared to that obtained by standard mechanical lysis and short read sequencing methods (see Methods). Although precise rank order relative abundances varied, we noted higher Shannon diversity from the present approach (P2: 2.0 vs. 1.14; P1: 2.0 vs. 1.8). We also detected all genera represented by more than 200 short reads from the traditional short read sequencing in the long read data.

Our assembly and post-processing workflow yielded whole-assembly N50 values of 453 kbp, 571 kbp and 564 kbp for the three samples P1, P2-A and P2-B. In comparison, the short read approach did not exceed assembly N50 of 34 kbp across samples P1 and P2-A, in spite of 3- to 6-fold more read data (37-38 Gbp). Our approach also surpassed the read cloud N50 values of 116 kbp and 12 kbp. However, read cloud and short read assemblies were between 1.5- and 2.1-fold larger than corresponding long read assemblies, likely due to the much greater volume of raw data available from these datasets (Supplementary Table 3).

Contigs from each approach were binned to form draft genomes, which were evaluated and assigned ‘High Quality’, ‘Complete’ and ‘Incomplete’ labels as described^9^. Briefly, drafts at least 90% complete and with at most 5% contamination are termed ‘Complete’, and drafts also containing at least one each of the 5S, 16S and 23S rRNA loci, as well as at least 18 tRNA loci, are labeled ‘High Quality’. All others are ‘Incomplete’. While read cloud and short read methods produced more complete bins and a comparable number of high quality bins compared to the long read approach, the long read approach produced bins with much higher contiguity (Figure 1). The present approach yielded nine high quality genomes with N50 over 2 Mbp, whereas the read cloud approach yielded only one. Short read bins never exceeded 550 kbp. Finally, the present approach yields a comparable quantity of high quality genomes at far higher contiguity with lower capital equipment requirement, sequencing cost and turnaround time (Supplementary Table 4).

**Figure 1.**
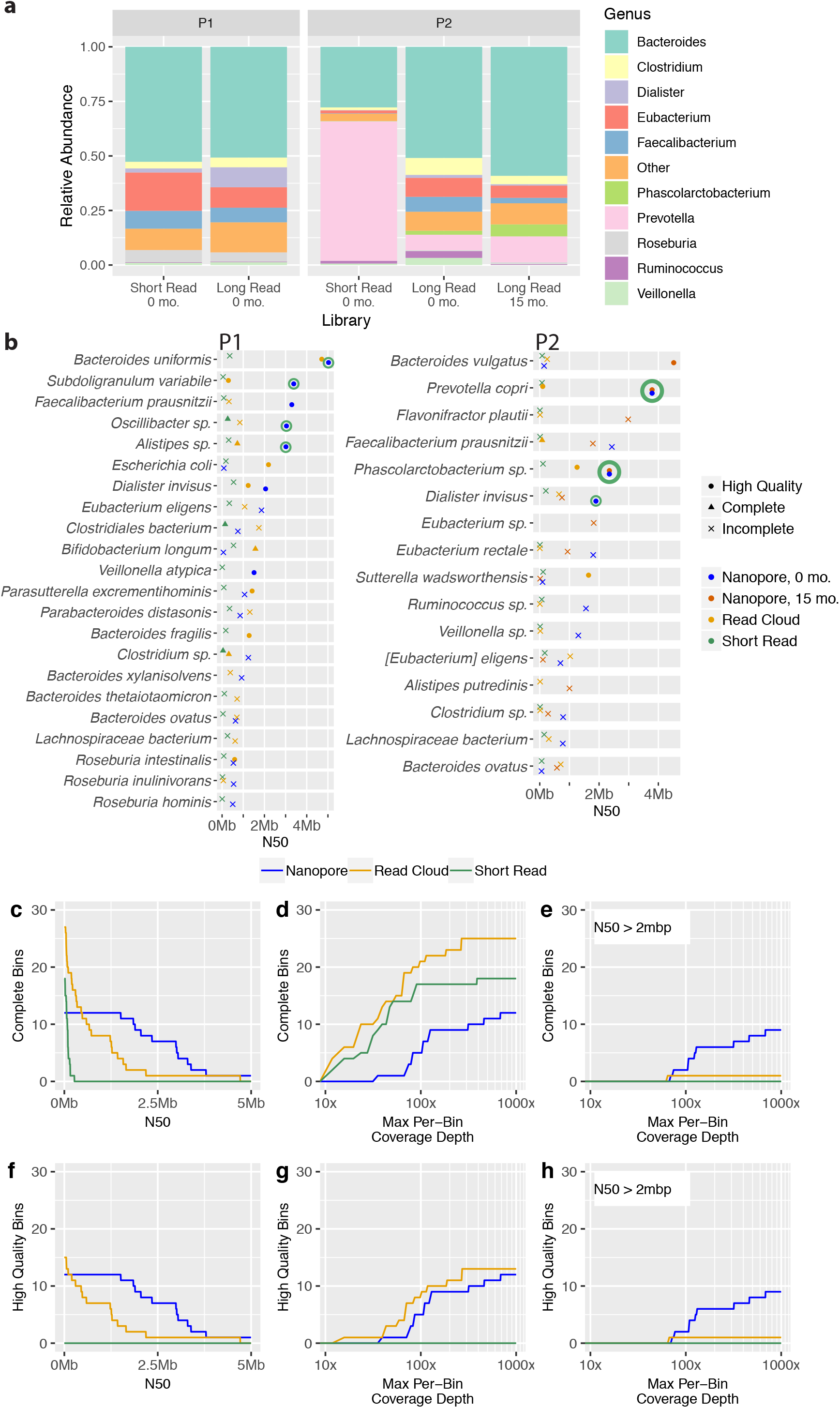
Taxonomic read composition and per-organism assembly contiguity for healthy gut assemblies, overall genome draft counts in two healthy human gut microbiomes (samples P1, P2-A). Nanopore sequencing and assembly (blue) demonstrates better assembly contiguity than read cloud (gold) and short read (green) approaches, but produced a smaller overall assembly with fewer complete drafts at the overall sequence coverage obtained. (a) Relative genus-level abundances are shown for a conventional workflow consisting of bead-beating extraction and short read sequencing, as well as the present workflow consisting of high molecular weight DNA extraction and long read sequencing. (b) For all organisms achieving assembly N50 of at least 500 kbp by any approach, genome draft quality and contiguity are shown for long reads, read clouds and short reads. Shapes indicate draft quality. Circularized genomes are indicated by green circles. (c) Complete genome bins with a minimum N50. (d) Complete genome bins below a given read coverage depth. Genome bins with lower read coverage originate from less abundant organisms. (e) Complete genome bins with N50 of > 2 Mbp below a given read coverage depth. (f) High quality genome bins with a minimum N50. (g) High quality genome bins below a given depth of read coverage. (h) High quality genome bins with an N50 exceeding 2 Mbp below a given read coverage depth.

Nanopore long read assembly yielded nine complete, circularizeable bacterial genomes across the three sequenced samples, and a maximum of four from a single sample (P1), compared to zero from the short read, read cloud, and synthetic long read approaches previously applied to these samples^9^. Assembled genomes are up to 5Mbp in length, and in several cases *(Prevotella copri, Subdoligranulum variabile, Phascolarctobacterium faecium,* and *Bacteroides uniformis)* represent the first closed genomes for their species. Closed genomes ranged in coverage depth between 75 *(Oscillibacter sp.)* and 785 *(P. copri).* Closed genomes were largely structurally concordant and similar in sequence to existing published genome sequences (Supplementary Figure 3, Supplementary Table 5), although in some cases we do note extensive strain divergence; for example, our closed *Dialister invisus* genome exhibits multiple large-scale inversions relative to the available reference (see below).

Completed bacterial genomes included two for *Prevotella copri* in samples P2-A and P2-B. This organism lacks a closed reference, in spite of extensive efforts to assemble *P. copri* and other members of the genus *Prevotella^10^.* Our previous efforts using read clouds, short reads and synthetic long reads to assemble these communities also had limited success with this organism, never exceeding a genome N50 of 130 kbp, in spite of coverage depth in excess of 4,800x^9^. The two *P. copri* genomes obtained from samples separated by 15 months display high concordance, with 99.94% of bases aligned and 99.89% nucleotide identity, suggesting nearly identical strain composition in the two time points.

The difficulty of assembling the *P. copri* genome stems from its high degree of sequence repetition. A direct assembly of highly abundant *k*-mers (*k*=101, occurring more than 5 times) found in our complete genome assembly yielded two insertion sequences (ISs) (see Methods), one 1.1 kbp IS66 family sequence and one 1.6 kbp IS1380 family sequence. These were found to be assembled in a total of 29 genomic loci between the two timepoints, but IS instances absent from the consensus assembly were detected directly in long reads at an additional 45 loci (see Methods). These insertion sites, whether fixed in the strain population or varying between strains, co-locate with breaks in short read and read cloud assemblies, illustrating their impact on these types of assembly (Figure 2).

**Figure 2.**
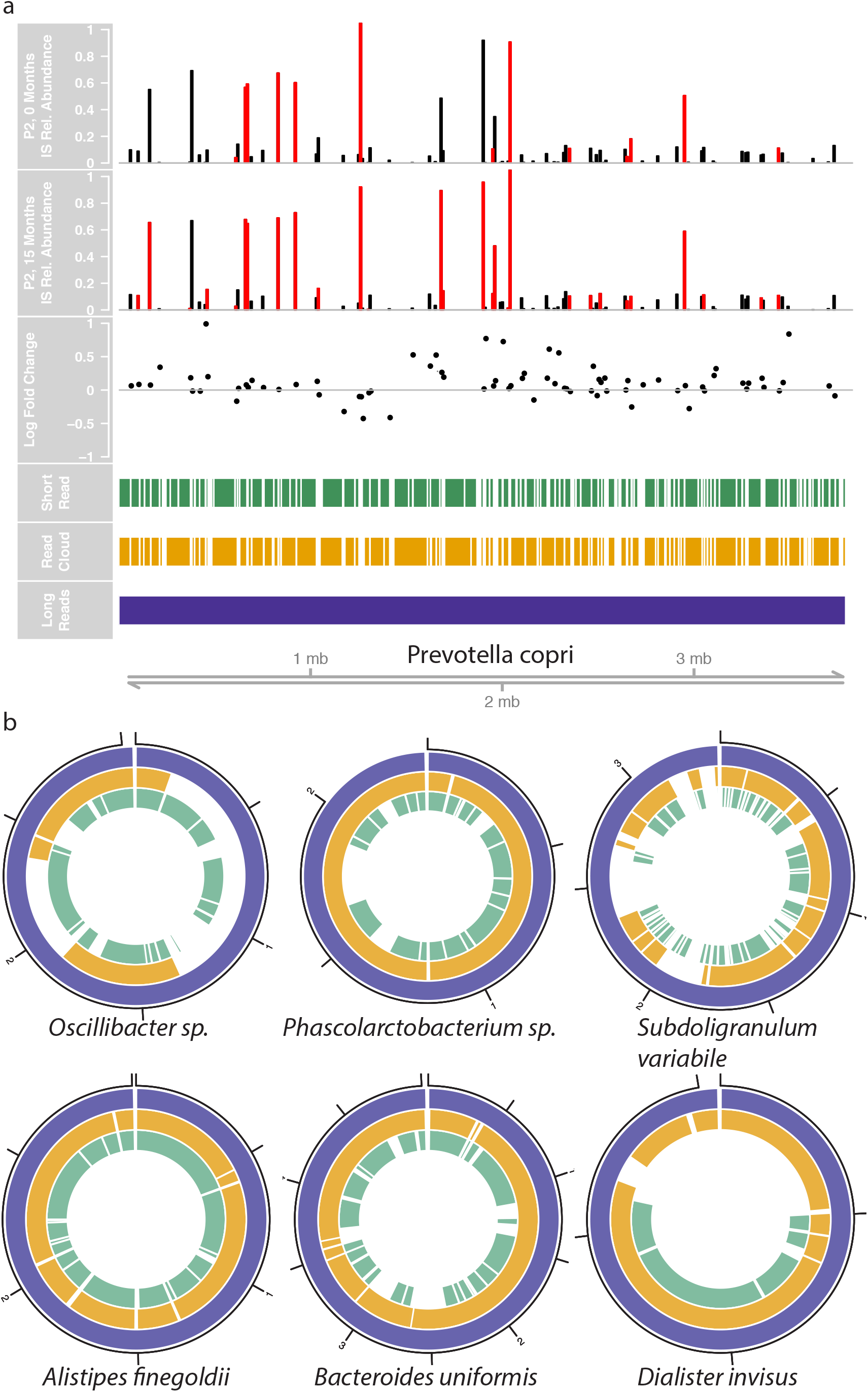
Genome assemblies, repeat structure and relative insertion sequence strain abundances of *Prevotella copri* and genome assembly comparisons for other closed genome assemblies. The *Prevotella copri* genome is difficult to assemble beyond insertion sequence sites due to their repetitiveness. For this reason, short read (green) and read cloud (gold) assemblies are highly fragmentary despite very high coverage (>4000× coverage depth). Long reads achieve a closed genome in spite of much lower coverage (318×) (blue). Relative abundances of strains carrying each insertion sequence instance are shown for 0-month and 15-month timepoints (first and second tracks), as well as log-fold change at each site between the two timepoints (third track). b) Finished genomes assembled by the present workflow (blue) are shown with corresponding bins obtained from read cloud (gold) and short read (green) approaches. Read cloud and short read approaches yield more fragmentary approaches, with large genomic regions missing due to incomplete binning.

Other complete genomes include *Phascolarctobacterium faecium* assembled in samples P2-A and P2-B at relative abundances of 4% and 1.41%, respectively. These assemblies are the first complete genomes for this species, and display high structural and nucleotide concordance with the closest available reference (Supplementary Figure 3; 98.9% identity, 88.5% sequence alignment) and with each other (99.81% identity, 99.97% sequence alignment). Sample P2-A also yielded the first circular genome for *Dialister invisus*, present in that sample at 1.03% relative abundance. We find similar structural divergence compared to the available reference (Supplementary Figure 3), and concordance with the read cloud draft, which contained identical large-scale structural inversions (99.90% identity, 99.97% sequence alignment) (Figure 2, Supplementary Table 5). Although the *Dialister invisus* assembled in sample P2-B was assessed complete, it was not found to be circularizable.

In sample P1, we obtained circular genomes for *Bacteroides uniformis* (6% abundance in long reads), *Alistipes finegoldii* (2% abundance), *Oscillibacter sp.* (0.14% abundance), and *Subdoligranulum variabile* (0.37% abundance). Of these, we were able to obtain structurally concordant reference sequences for all but *Subdoligranulum,* for which we could not locate a reference with more than 19% aligned bases, suggesting the possibility of a novel strain. Seven 16S rRNA loci were assembled in this genome, all bearing 98% sequence identity to the closest match from *Subdoligranulum variabile* strain BI114, for which no genome reference is available for comparison. Identity with read cloud assemblies in all cases was over 99.7%, with over 98.3% of bases aligning to the assembled draft (Supplementary Table 5). For all closed genomes, read cloud and short read assembly yielded more fragmentary assemblies, which were only partially recovered by binning (Figure 2).

Our approach relies on consensus refinement based on short read data to correct homopolymer errors intrinsic to the current nanopore sequencing technology. Although long read-based consensus refinement is possible and partially effective, we find that it cannot fully replace short read correction (Supplementary Figure 4). We found that uncorrected long read assembly demonstrated a 3% error rate with 3-mer homopolymers, assembled too short by an average of 0.5 nucleotides. This worsens to a 65% error rate on 6-mer homopolymers, which were assembled too short by an average of 1.3 nucleotides. On average, 63 homopolymers of length 3 or greater were found per kilobase of assembled sequence, of which 4.5 (7.1%) were found to require correction with short reads. CheckM, a tool which annotates genome completeness based on single copy core gene detection, demonstrates a low detection rate on uncorrected assemblies, consequently under-reporting genome completeness even on circularizeable whole-genome contigs. For instance, the genome annotated *Oscillibacter sp.* in sample P1 is annotated 38% complete in the uncorrected assembly. This rises to 68% complete after correction with long reads. With short read correction, the genome receives a 96% completeness annotation, compared to 98% for the sole available closed genome reference sequence (strain PEA192). The present workflow for sequencing and assembly can operate solely with long reads and will yield structurally correct and complete genomes, although with reduced nucleotide accuracy. Future advances in nanopore sequencing technology that decrease the homopolymer-repeat related errors will likely lessen or remove the requirement for supplemental short read sequencing to achieve genomes with high nucleotide fidelity.

Although the present approach has achieved effective assembly of bacterial genomes from metagenomes, we anticipate that future advances in metagenomic DNA extraction methods and nanopore long read assembly will improve read length and reduce the read coverage required to close genomes. In addition, epigenetic modification detection will add to future metagenomic studies by revealing phage and bacterial sequence methylation patterns, methylation-based contig binning approaches, and epigenetic regulation of bacterial DNA-protein interactions.

In conclusion, our approach assembles the first complete genome of *Prevotella copri,* an organism with high prevalence in non-western guts and with emerging, potentially strain-specific links to human health and disease^11,12^. The high copy number of IS66 and IS1380 family insertion sequences in this genome limit the effectiveness of short read approaches, despite receiving over 4,800x coverage in an earlier metagenomic sequencing study^9^ and extensive isolate sequencing in a separate effort^10^. IS1380 has been previously reported to carry an outward-facing promoter capable of upregulating adjacent gene sequences, and has been found to impact antibiotic resistance gene regulation and resistance phenotype^13^. We anticipate that this approach will help illuminate the role of repetitive classes of genomic elements with important effects on cellular and clinical phenotypes, and facilitate efforts to broaden human microbiome research to global populations where *Prevotella* are highly prevalent^14^. Closing these and other genomes will allow investigation into the complete functional repertoire and potential phenotypes of individual microbes, even when these organisms are difficult to culture or are found in mixed communities, facilitating future research in important human microbiomes and poorly characterized microbial communities such as soil and marine sediment.

## Supporting information

## Methods

### DNA extraction

Short read and read cloud libraries were prepared as previously described^9^. Previously, DNA was extracted from samples P1 and P2-A with a commercial extraction kit using bead-beating lysis.

For high molecular weight (HMW) extraction, approximately 0.7g frozen stool was aliquoted into 2mL eppendorf tubes (Eppendorf, Hamburg, Germany) with a 4mm biopsy punch (Integra Miltex, Plainsboro, NJ) and suspended in 500μL PBS (Fisher Scientific, Waltham, MA) with brief gentle vortexing. 5uL of lytic enzyme solution (Qiagen, Hilden, Germany) was added and the samples were mixed by gentle inversion six times, then incubated for one hour at 37°C. 12μL 20% (w/v) SDS (Fisher Scientific, Waltham, MA) was added with approximately 100μL vacuum grease (Dow-Corning, Midland, MI) functioning as phase lock gel^15^. 500μL phenol chloroform isoamyl alcohol at pH 8 (Fisher Scientific, Waltham, MA) was added, and samples were gently vortexed for five seconds, then centrifuged at 10,000g for five minutes with Legend Micro 21 microcentrifuge (Fisher Scientific, Waltham, MA). The aqueous phase was then decanted into a new 2mL tube.

Next, DNA was precipitated with 90μL 3M sodium acetate (Fisher Scientific) and 500uL isopropanol (Fisher Scientific) for ten minutes at room temperature. After inverting three times slowly, samples were incubated at room temperature for 10 minutes, then centrifuged 10 minutes at 10,000g. The supernatant was removed and the pellet was washed two times with freshly prepared 80% (v/v) ethanol (Fisher Scientific). The pellet was then air dried with heating for ten minutes at 37°C or until the pellet was matte in appearance, and then resuspended in 100μL nuclease-free water (Ambion, Thermo Fisher Scientific, Waltham, MA). 1mL Qiagen buffer G2, 4μL Qiagen RNase A at 100mg/mL, and 25μL Qiagen Proteinase K were added, the samples were then gently inverted three times, and then were incubated 90 minutes at 56°C. After the first 30 minutes, pellets were dislodged by a single gentle inversion.

One Qiagen Genomic-tip 20/G column per sample was equilibrated with 1mL Qiagen buffer QBT and allowed to empty by gravity flow. Samples were gently inverted twice, applied to columns and allowed to flow through. Three stool extractions were combined per column. Columns were then washed with 3mL Qiagen buffer QC, then DNA was eluted with 1mL Qiagen buffer QF prewarmed to 56°C. Eluted DNA was then precipitated by addition of 700μL isopropanol followed by inversion and centrifugation for 15 minutes at 10,000g. The supernatant was carefully removed by pipette, and pellets were washed with 1mL 80% (v/v) ethanol. Residual ethanol was removed by air drying ten minutes at 37°C. This was followed by resuspension of the pellet in 100μL water overnight at 4°C without agitation or any kind.

DNA was then size selected with a modified SPRI bead protocol as described ^16^, with minor modifications: beads were added at 0.9×, and eluted DNA was resuspended in 50μL water. The concentration, purity and fragment size distribution of extracted DNA was then quantified with the Qubit fluorometer (Thermo Fisher Scientific, Waltham, MA), Nanodrop (Thermo Fisher Scientific), and Tapestation 2200 (Agilent Technologies, Santa Clara, CA), respectively (Supplementary Table 1).

### Sequencing

Extracted DNA samples were prepared for long read sequencing with the Oxford Nanopore Technologies (ONT, Oxford, UK) Ligation library preparation kit according to the manufacturer’s standard protocol. Libraries were sequenced with the ONT MinION sequencer using rev C R9.4 flow cells, allocating one flowcell per sample. The sequencer was controlled with the MinKNOW v2.2.12 software running on a MacBook Pro (model A1502, Apple, Cupertino, CA), with data stored to a Vectotech 2Tb SSD hard drive. Sequencing runs were scheduled for 48 hours, and allowed to run until fewer than 10 pores remained functional. After sequencing, data were uploaded to the Stanford Center for Genomics computational cluster for analysis (see below). Short read libraries were prepared and sequenced as described previously^9^.

### Sequence assembly and analysis

Raw data were basecalled with Albacore v2.3.1, and assembled in two separate runs with Canu v1.7.1 with the -nanopore preset parameter ^17^. The two runs differed by the estimated genomeSize parameter, provided as either 50m or 100m. The two separate assemblies were then merged with quickmerge v0.40^18^, circularized with Circlator v1.5.5^19^ and Encircle (present study, see below), and then polished with either Medaka ^20^ or a parallelized version of Pilon v1.22 ^21^ (present study) for long read or short read consensus refinement, respectively. In order to parallelize Pilon, reference sequences were divided into 100kb segments, short reads aligning to each segment were downsampled to at most 40x coverage depth, and Pilon was used to detect errors within the reference and read subset. These errors were then aggregated across all subset runs and used to generate a refined consensus with bcftools^22^. Errors found in homopolymers were identified with an in-house script, homopolymer_error_analyzer. Sequences are binned and annotated as previously described ^9^.

There is presently no straightforward, comprehensive method for determining circularity in metagenome-assembled genomes. A minority of the circular genomes we obtained *(D. invisus* in P2-A, *P. copri and Phascolarctobacterium* in P2-B) were circularized by an existing genome circularization tool. In several cases, assembled genome contigs extended beyond the wrap-around point of the circular chromosome, resulting in what we term over-circularization (supp fig 5). Over-circularized contigs contain redundant sequences at their termini which spuriously increase apparent contamination when assessed by CheckM. In order to trim over-circularized contig ends in order to obtain a nonredundant, circular genome, we developed Encircle, a utility which performs contig self-alignment with Mummer^23^ and determines when over-circularization has taken place, then outputs precise trim coordinates to circularize the genome. The genomes of *P. copri, Phascolarctobacterium sp.,* and *Dialister invisus* in sample P2-A, as well as *Oscillibacter sp.* and *Subdoligranulum* (supp figure 5) in sample P1, were overcircularized and required trimming. In addition, the genomes of *Bacteroides uniformis* and *Alistipes finegoldii* were determined to be circular by concatenating the first and last 20kbp of the assembled genome, mapping long reads to the junction, and inspecting alignments for reads spanning the gap; *B. uniformis* was found to be slightly overcircularized by 10kbp (below the limit of detection of Encircle), and *A. finegoldii* was found to be perfectly circularizeable.

Binning was performed and evaluated as previously described^9^. Due to the complete genomes present in our assemblies, binning became unnecessary for some organisms, and instead led to several cases of genomic contamination as assessed with CheckM. In cases where >5% contamination occurred in a bin with one genome-scale contig and several much smaller (<100kbp) sequences, the smaller sequences were removed and the largest sequence was re-evaluated with CheckM^24^, in two cases yielding complete and uncontaminated genomes.

Long and short reads were taxonomically classified with Kraken^25^, and Shannon diversity was calculated with vegan^26^. Figures were generated with ggplot2^27^, gviz^28^, doBy^29^ and reshape2^30^. All workflows were implemented with Snakemake^31^.

### Insertion sequence strain diversity

K-mers represented more than 6 times in the *Prevotella copri* assemblies were identified with Jellyfish2^32^. These were assembled with SPAdes^33^ two obtain two full-length insertion sequences. These sequences were located in the genome assemblies by alignment with minimap2^34^. In order to locate additional unassembled insertion sequences present in strains of *P. copri,* reads containing insertion sequences were identified by alignment with minimap2, then 200 bases immediately upstream of the insertion sequence were taken from each read and aligned to the genome assembly.

In order to quantify the relative abundance of *P. copri* strains carrying each IS instance, long reads were first aligned to the assembled IS sequences. Long reads containing IS sequences were isolated, and flanking sequences 200bp upstream of the IS were extracted and realigned to the genome assembly. IS-flanking sequence depth was compared to local overall coverage depth to obtain the relative abundance of strains carrying a given IS. Only 18 insertion sites carried fixed ISs and a further 56 sites showed a mixture of strains with and without an IS (Figure 2).

### Data availability

All sequence data, whole metagenome assemblies and individual completed genomes can be found at NCBI BioProject under accession PRJNA508395.

### Code availability

All workflows and associated environments and tools can be found at https://github.com/elimoss/metagenomics_workflows/.

## Acknowledgements

The authors would like to acknowledge Gavin Sherlock and the members of the Bhatt lab for helpful advice and assistance. E.L.M. was supported by National Science Foundation Graduate Research Fellowship DGE-114747. This work was supported by the Damon Runyon Clinical investigator award to ASB. Computational work was supported by NIH S10 Shared Instrumentation Grant 1S10OD02014101.

## Supplementary Figure Legends

<u>Supplementary Figure 1</u> Overview of the molecular and informatic workflow steps. a) Extraction consists of enzymatic degradation of bacterial cell walls followed by an initial DNA extraction in phenol-chloroform. This is followed by a proteinase K and RNase A digestion at high temperature and purification with a gravity column. Finally, small fragments are removed by modified SPRI bead size selection. b) After sequencing and basecalling, read sequences are assembled twice with varying genomeSize parameter values. These two assemblies are merged, then circular sequences are identified and trimmed. The consensus sequence is refined by either short-read or long-read polishing.

<u>Supplementary Figure 2</u>

Histogram of total bases versus read length for the three samples sequenced with the current approach. Read lengths vary between <1kbp to >100kbp, with N50 values between 5kbp and 10kbp.

<u>Supplementary Figure 3</u>

Reference alignment dotplots for closed genomes obtained by nanopore long read sequencing and assembly. Although assemblies share broad structural similarity to available references, there are cases where observed organisms are significantly structurally diverged (e.g. *Dialister*) and in one case bears minimal similarity to the available reference (*Subdoligranulum*).

<u>Supplementary Figure 4</u>

Homopolymer count as a function of length, and homopolymer error in assembled sequence as a function of length in corrected sequence. We found that uncorrected long read assembly demonstrated a 3% error rate with 3-mer homopolymers, assembled too short by an average of 0.5 nucleotides. This worsens to a 65% error rate on 6-mer homopolymers, which were assembled too short by an average of 1.3 nucleotides. On average, 63 homopolymers of length 3 or greater were found per kilobase of assembled sequence, of which 4.5 (7.1%) were found to require correction with short reads.

<u>Supplementary Figure 5</u>

Nanopore long read assembly in some cases produces over-circularized genomes. These are sequences that are assembled beyond the wrap-around point, resulting in (a) redundant sequence which are detected and trimmed with the Encircle utility (present study). These sequences can be visualized as (b) corner-cutting off-diagonal alignments within contig self-alignment dotplots, such as that shown for the untrimmed *Subdoligranulum variabile* assembly.

