## Supplementary figures and images for "Generating closed bacterial genomes from long-read nanopore sequencing of microbiomes"

### Supplementary file 1

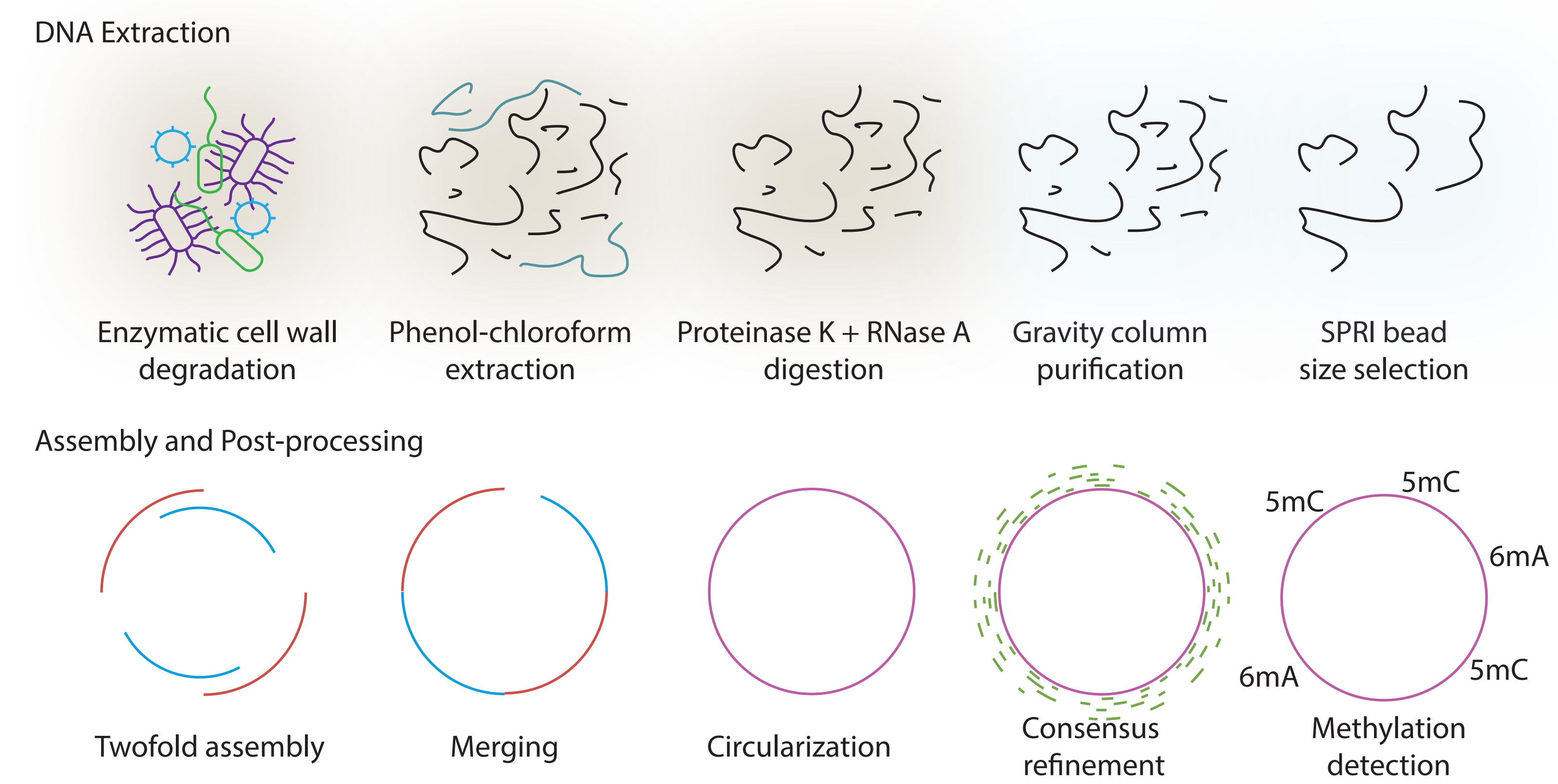

### Supplementary file 2

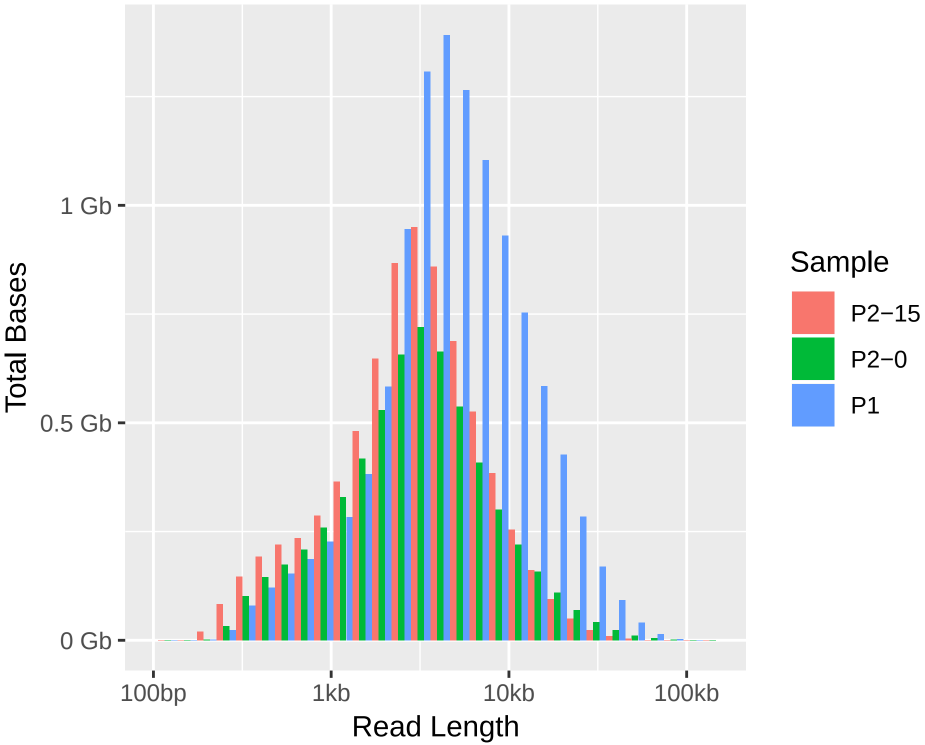

### Supplementary file 3

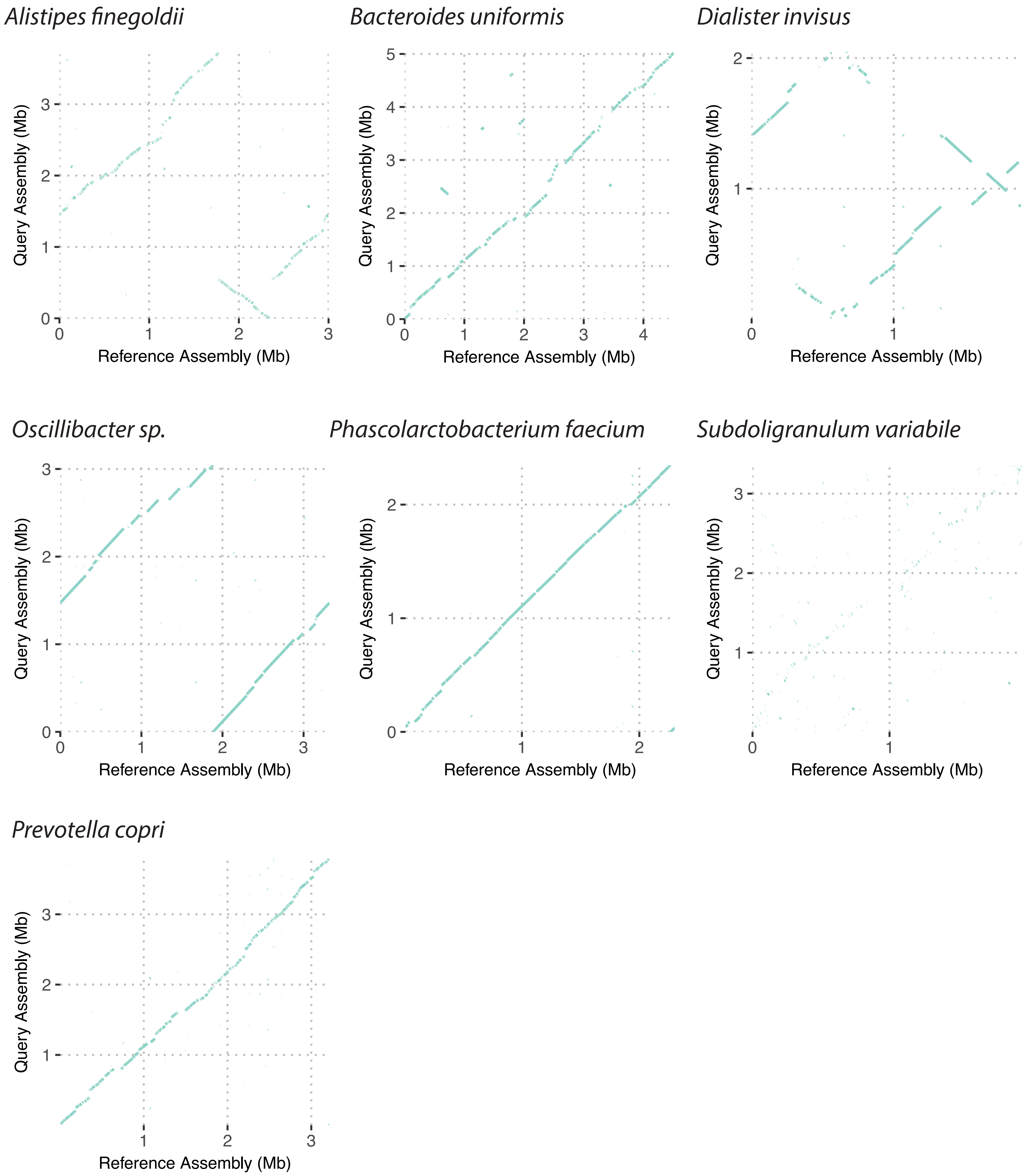

### Supplementary file 4

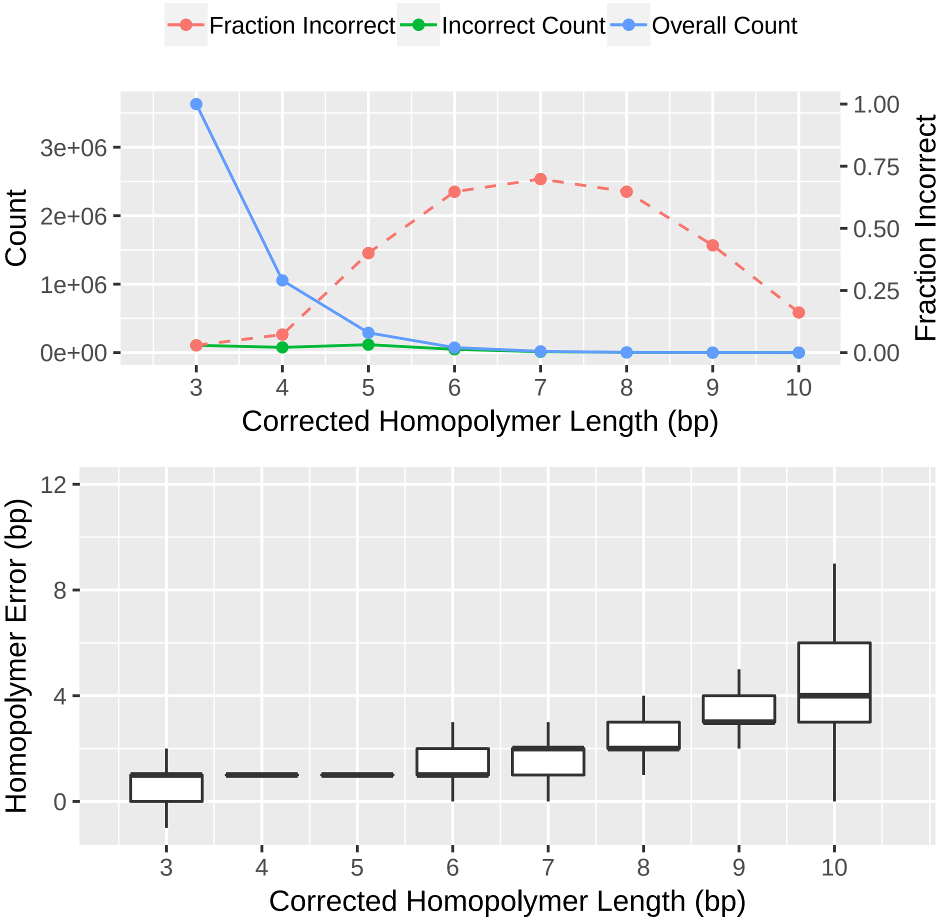

### Supplementary file 5

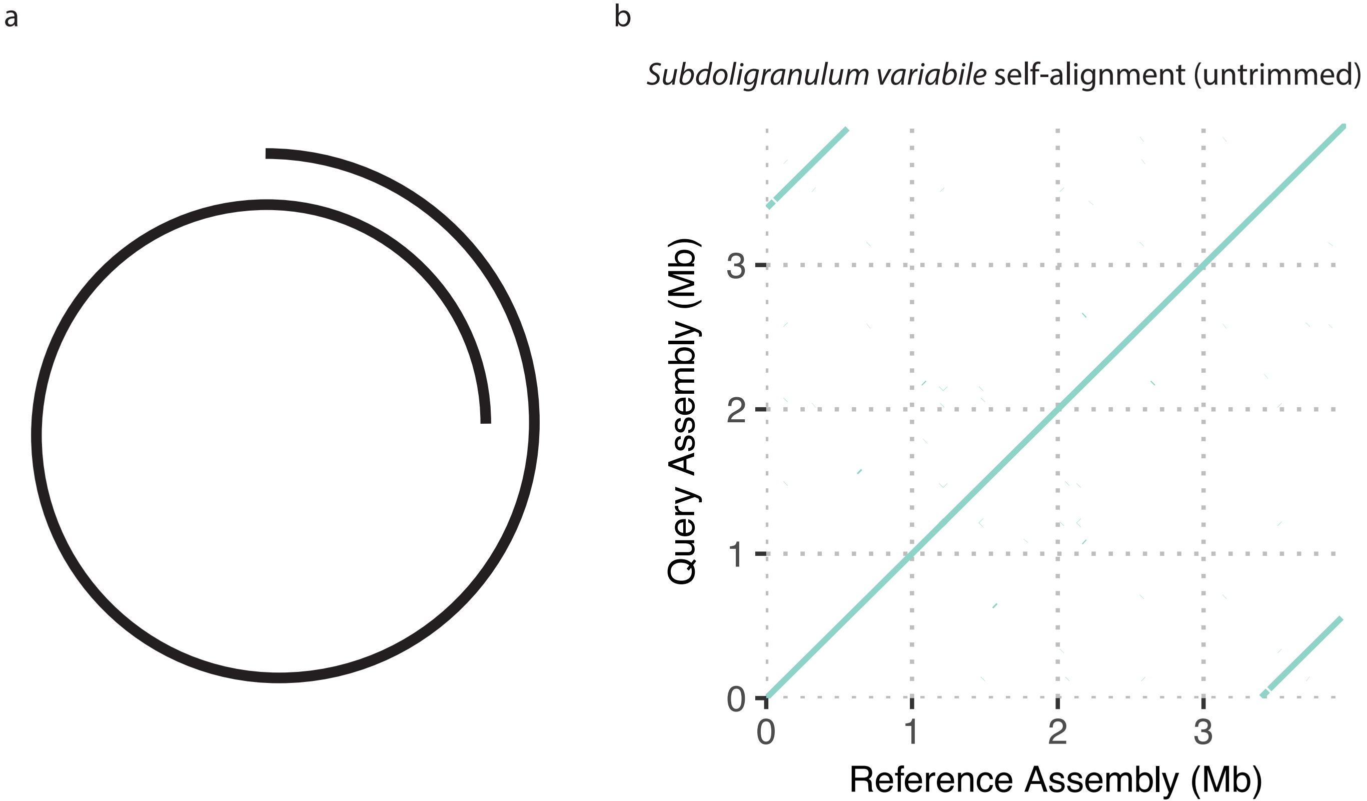
